# Genomic and Phenotypic Comparison of Polyhydroxyalkanoates Producing Strains of genus *Caldimonas/Schlegelella*

**DOI:** 10.1101/2023.09.27.559687

**Authors:** Jana Musilova, Xenie Kourilova, Kristyna Hermankova, Matej Bezdicek, Anastasiia Ieremenko, Pavel Dvorak, Stanislav Obruca, Karel Sedlar

## Abstract

Polyhydroxyalkanoates (PHAs) have emerged as an ecologically friendly alternative to conventional polyesters. In this study, we present a comprehensive analysis of the genomic and phenotypic characteristics of three non-model thermophilic bacteria known for their ability to produce PHAs: *Schlegelella aquatica* LMG 23380^T^, *Caldimonas thermodepolymerans* DSM 15264, and *C. thermodepolymerans* LMG 21645 accompanied by a comparison with the type strain *C. thermodepolymerans* DSM 15344^T^. We have assembled the first complete genomes of these three bacteria and performed the structural and functional annotation. This analysis has provided valuable insights into the biosynthesis of PHAs and has allowed us to propose a comprehensive scheme for the carbohydrate metabolism in the studied bacteria. Through phylogenomic analysis, we have confirmed the synonymity between *Caldimonas* and *Schlegelella* genera, and further demonstrated that *S. aquatica* and *S. koreensis*, currently classified as orphan species, belong to the *Caldimonas* genus.

**Summary:** The genomic and phenotypic analysis of *Schlegelella aquatica* LMG 23380^T^ and *Caldimonas thermodepolymerans* DSM 15264 and LMG 21645 sheds light on the production of sustainable polyesters known as polyhydroxyalkanoates (PHAs). The genome assembly and functional annotation highlight key genes related to PHA production and other important traits. Notably, *C. thermodepolymerans* stands out with its unique *xyl* operon, making it a highly promising candidate for biotechnological PHA production from xylose-rich lignocellulosic resources. The study emphasizes the importance of a polyphasic approach combining genotypic and phenotypic analyses in prokaryotic taxonomy, emphasizing the need for exploration in the genomic era. By uncovering the key traits of these bacteria, this research opens new horizons towards sustainable production of environmentally friendly polyesters.

## Introduction

Polyhydroxyalkanoates (PHAs) are polyesters accumulated by numerous prokaryotes in the form of intracellular granules to serve as carbon and energy storage materials and to enhance stress robustness of bacterial cells (Obruca et al. 2018). Moreover, PHAs have emerged as environmentally friendly substitutes for petroleum-based polymers, offering a sustainable, renewable, biodegradable, compostable and also biocompatible alternative (Koller et al. 2017; Sabapathy et al. 2020). Although the production of bioplastics is seen as the way of the future and an integral part of the circular economy, less than 1% of total plastics production comes from the bioplastics industry (Shogren et al. 2019).

Extremophiles are organisms able to survive and even prosper in extreme conditions such as acidic or basic pH levels, the presence of toxic elements, and immoderate temperatures (Rothschild and Mancinelli 2001; Martin and McMinn 2018). Since the cultivation of extremophiles can be processed in a semi-sterile or even unsterile mode, which significantly lowers the cost of biotechnological procedures, these organisms have gained a key role in the so-called “Next-generation Industrial Biotechnology” (NGIB) concept (Chen and Jiang 2018) in recent years (Coker 2016). Further benefits come from the employment of thermophiles, bacteria that thrive at temperatures above 45°C. Although biotechnological processes operate at high temperatures, they can be energetically feasible because the metabolic heat and energy released during mixing can be used to heat the bioreactor. In addition, cooling costs are modest because ambient air can be used to cool the process (Ibrahim and Steinbüchel 2010; Obruča et al. 2022).

Several potent thermophilic PHAs producers can be found in the recently proposed family *Sphaerotilaceae*, which, after the taxonomic revisions based on phylogenomic comparisons, contains several genera, including the genus *Caldimonas* (Liu et al. 2022). The most promising group of organisms within this genus is formed by species belonging originally to the genus *Schlegelella* (Nunes et al. 2021) with the type species Schlegelella *thermodepolymerans*, which was initially studied for its ability to degrade PHAs such as 3-hydroxybutyrate and 3-mercaptopropionate copolymers (Elbanna et al. 2003). Moreover, an extraordinary ability of the type strain *S. thermodepolymerans* DSM 15344^T^ to utilize xylose and synthesize 3-hydroxybuytrate and 3-hydroxyvalerate copolymers was recently reported by our group (Kourilova et al. 2020) and the bacterium was identified as a promising candidate for industrial production of PHAs from various xylose rich lignocellulose-based resources (Kourilova et al. 2021). Subsequently, we provided the first complete genome assembly and the functional annotation (Musilova et al. 2021). With the availability of complete genome sequences, *Schlegelella thermodepolymerans* was found to be a homotypic synonym of *Caldimonas thermodepolymerans* (Liu et al. 2022). Since the genomes of the two other *Schlegelella* species were not available and 16S rRNA gene phylogeny was found to be insufficient to infer evolutionary relationships in the family *Sphaerotilaceae*, the two orphaned species, *Schlegelella aquatica* and *Schlegelella koreensis*, remained in the genus *Schlegelella*. However, additional research is needed to gain a comprehensive understanding of the evolutionary relationships within the genus, as well as to uncover the full genomic, metabolic, and biotechnological potential of these non-model bacteria.

Here, we compared the newly assembled genome of the type strain of S. aquatica LMG 23380^T^ with the recently published genome of the type strain of *S. koreensis* ID0723^T^ and other genomes from the genus *Caldimonas* including two newly assembled genomes of non-type *Caldimonas thermodepolymerans*, formerly *Schlegelella thermodepolymerans*, species. Besides further revisiting their taxonomy, we aimed at their genotypic and phenotypic comparison because all *Caldimonas /Schlegelella* species remain underexplored, which prevents their possible use in industrial biotechnology. Apart from comparing their genomes, examining the ability to utilize various carbon sources, and exploring their antibiotic susceptibility, we also identified genes coding the key enzymes in core carbohydrate and PHA metabolism and proposed a basic metabolism scheme.

### Experimental Procedures

#### Growth conditions and phenotypic analysis

In the experiments, four different strains of the genus *Caldimonas /Schlegelella* were used. Two of them were purchased in Leibnitz Institute DSMZ German Collection of Microorganisms and Cell Cultures, Braunschweig, Germany: *Caldimonas thermodepolymerans* DSM 15344^T^ and *Caldimonas thermodepolymerans* DSM 15264. The other two were obtained in BCCM/LMG Bacteria Collection in the Laboratory of Microbiology, Department of Biochemistry and Microbiology, Faculty of Sciences of Ghent University: *Schlegelella aquatica* LMG 23380^T^ and *Caldimonas thermodepolymerans* LMG 21645. The bacteria were cultivated in two steps. In the first part of the cultivation, a nutritionally rich medium (Nutrient broth) was used (100 mL Erlenmeyer flasks, filling 50%). This phase entailed 20 hours at 50 °C and constant shaking at 180 rpm. Subsequently, for the production phase, mineral medium was used (9.0 g/L Na_2_HPO_4_ · 12 H_2_O, 1.5 g/L KH_2_PO_4_, 1.0 g/L NH_4_Cl, 0.2 g/L MgSO_4_ · 7 H_2_O, 0.02 g/L CaCl_2_ · 2 H_2_ O, 0.0012 g/L Fe^(III)^NH_4_ citrate, 0.5 g/L yeast extract) with 20 g/L carbon sources (cellobiose, glucose, xylose) and trace element solution (50.0 g/L EDTA, 13.8 g/L FeCl_3_ · 6 H_2_O, 0.84 g/L ZnCl_2_, 0.13 g/L CuCl_2_ · 2 H_2_O, 0.1 g/L CoCl_2_ · 6 H_2_O, 0.016 g/L MnCl_2_ · 6 H_2_O, 0.1 g/L H_3_BO_3_, dissolved in distilled water) and 5 % v/v of culture from complex medium (250 mL Erlenmeyer flasks, filling 40%). This part of the cultivation took 72 hours, also at 50 °C and constant shaking at 180 rpm.

Duplicates of biomass samples (2 × 10 mL) were determined gravimetrically. The biomass was centrifuged (6 000 × g, 5 min), then washed with 10 mL distilled water, and centrifuged again. The supernatant was discarded and the pellet of biomass was dried to constant weight. PHA content in dry biomass was determined by gas chromatography with a flame ionization detector as described previously (Obruca et al. 2014).

#### Antibiotic susceptibility testing

The sensitivity of studied *Caldimonas /Schlegelella* strains to kanamycin (Serva, Germany), streptomycin, (Serva, Germany), gentamycin (Roth, Germany), tetracycline (Roth, Germany) and chloramphenicol (Serva, Germany) was examined. Cell cultures inoculated from overnight cultures (these were prepared in 20 mL NB medium in 100 mL Erlen flasks inoculated directly from glycerol stocks) to the starting OD_600_ of 0.05 were grown in 3 mL of NB medium (Himedia, India) in 15 mL plastic tube at 50°C with shaking (200 rpm, Biosan ES-20/60) for 72 h. With all of the antibiotics except streptomycin, ten concentrations were tested – 0, 2.5, 5, 7, 12.5, 25, 50, 100, 150, and 200 μg/mL. Three additional concentrations of 300, 400, and 500 μg/mL were checked with regards to streptomycin. All the experiments were performed in two biological replicates. Raw OD_600_ data from all of the measurements is provided in Supplementary File 2.

#### DNA extraction and sequencing

High molecular weight genomic DNA for long-read sequencing was extracted using MagAttract HMW DNA kit (Qiagen, NL) in accordance with the manufacturer’s instructions. The DNA purity was checked using NanoDrop (Thermo Fisher Scientific, USA), the concentration was measured using Qubit 4.0 Fluorometer (Thermo Fisher Scientific, USA), and the length was measured using Agilent 4200 TapeStation (Agilent Technologies, USA). The Ligation Sequencing 1D Kit (Oxford Nanopore Technologies, UK) was used for libraries preparation and the sequencing was performed using R9.4.1 Flow Cell on the Oxford Nanopore Technologies (ONT) MinION platform. Genomic DNA for the high-throughput short-read sequencing was extracted using GenElut Bacterial Genomic DNA Kit (Sigma-Aldrich, USA) in accordance with the manufacturer’s instructions. Sequencing libraries were prepared using KAPA HyperPlus kit and sequencing was carried out using Miseq Reagent kit v2 (500 cycles) on the MiSeq platform (Illumina, USA).

#### Genome assembly

The assemblies were completed in two steps. First, ONT reads were assembled into the initial sequence, and afterward, Illumina reads were mapped onto the initial sequence to produce hybrid assemblies. In the first step, the ONT reads were initially basecalled using Guppy v3.4.4, and the reads were assembled using Flye v2.8.1 (Kolmogorov et al. 2019). Quality check was done using MinIONQC (Lanfear et al. 2019). The assembly was polished with Racon v1.4.20 (Vaser et al. 2017) and Medaka; PAF files were generated using minimap2 v2.24 (Li 2018). In the second step, Illumina paired end (PE) reads were firstly quality-checked using FastQC v0.11.5 and MultiQC v1.7 (Ewels et al. 2016) and, subsequently, adapter and the quality trimming of the Illumina reads were performed using Trimomatic v0.36 (Bolger et al. 2014). The trimmed reads were mapped on the Nanopore initial assembly using BWA v07.17 (Hi 2013). The polishing was done using Pilon v1.24 (Walker et al. 2014) and BAM files were generated using SAMtools v1.14 (Li et al. 2009). Finally, the sequences were rearranged according to the origin of replication (oriC) in order for DnaA to be the first gene on the sense strand.

#### Genome annotation

The genome structural annotation was completed through the NCBI Prokaryotic Genome Annotation Pipeline (PGAP) (Tatusova et al. 2016). Operons prediction was performed using Operon-mapper (Taboada et al. 2018), and the results were further processed to obtain polycistronic operons. The genes involved in carbohydrate metabolism were primarily identified through the search for homologous sequences using the NCBI BLAST tool (Altschul et al. 1990), along with a comprehensive literature search. For the genes associated with PHA metabolism, the PHA Depolymerase Engineering Database (Knoll et al. 2009) was employed in addition to the aforementioned methods.

Functional annotation of protein-coding genes was performed with eggNOG-mapper (Cantalapiedra et al. 2021), Operon-mapper (Taboada et al. 2018), and batch CD-Search (Marchler-Bauer and Bryant 2004) by classifying them into clusters of orthologous groups (COGs) and assigning them particular functional categories. Outputs from the abovementioned tools were combined into the consensus result using COGtools (available from https://github.com/xpolak37/COGtools) and visualized in barplots for comparison of particular genomes. Methylated bases were detected from ONT raw data by nanopolish (Simpson et al. 2017); methylation motifs were inferred using STREME Command-Line version (Bailey 2021). The detection of Restriction-Modification (R-M) systems was done using the Restriction Enzyme Database’s (REBASE) tools (Roberts et al. 2015) and KEGG Database (Kanehisa 2000). Antibiotic-resistant genes search was performed using Resistance Gene Identifier (RGI) 6.0.0 included in the Comprehensive Antibiotic Resistance Database (CARD) 3.2.5 (Alcock et al. 2020).

Clustered Regularly Interspaced Short Palindromic Repeats (CRISPR) arrays identification was carried out via the CRISPRDetect tool (Biswas et al. 2016).

Digital DNA to DNA hybridization (dDDH) values were calculated using the Type (Strain) Genome Server (TYGS) (Meier-Kolthoff and Göker 2019). A phylogenomic tree was constructed with a bootstrapping method using PhyloPhlAn 3.0.60 (Segata et al. 2013) and its internal database of approximately 400 genes conserved across the bacterial domain.

## Results

### Genome assembly & structural annotation

*Schlegelella aquatica* LMG 23380^T^ and *Caldimonas thermodepolymerans* (formerly *Schlegelella thermodepolymerans)* DSM 15264 and LMG 21645 genomes were assembled using a hybrid approach with initial coverage of 261×, 1,440×, and 160×, respectively. Over 44 thousand Oxford Nanopore Technologies (ONT) reads with a median read length of 3.44 kb were used to generate an initial assembly of S. aquatica LMG 23380^T^. Over 1.6 million high-quality (average Phred score Qlll≈lll34) Illumina reads (97% of all Illumina reads) were subsequently mapped to the initial assembly. *C. thermodepolymerans* DSM 15264 was reconstructed from over 901 thousand ONT reads with a median read length of 2.51 kb and more than 2.67 million high-quality (average Phred score Qlll≈lll36) Illumina reads (86% of all Illumina reads) mapped to the initial assembly. *C. thermodepolymerans* LMG 21645 initial assembly was reconstructed from over 14 thousand ONT reads with a median read length of 2.7 kb. Subsequently, more than 1.8 million high-quality (average Phred score Qlll≈lll36) Illumina reads (89% of all Illumina reads) were mapped to the initial assembly. The whole process resulted in circular sequences with coverage of 194×, 1,324× and 85×, corresponding to S. aquatica LMG 23380^T^, *C. thermodepolymerans* DSM 15264, and *C. thermodepolymerans* LMG 21645, respectively. The final assemblies have been deposited at the DDBJ/EMBL/GenBank under accession numbers CP110257, CP110416, and CP110415.

The genome lengths range from 3.3 to 4.0 Mbp and contain from 3,069 to 3,856 open reading frames (ORFs). The majority of all ORFs consist of protein-coding sequences (CDSs), but several pseudogenes were also identified. Table 1 summarizes the main features of the newly assembled genomes along with the genome of the type strain *C. thermodepolymerans* DSM 15344^T^ (Musilova et al. 2021), available in the GenBank NCBI database under the accession CP064338.

**Table 1.** Chromosomal features of *Schlegelella aquatica* (Sa) LMG 23880^T^, *Caldimonas* thermodepolymerans (Ct) DSM 15264 and LMG 21645, and type strain DSM 15344^T^.

| Feature | *Sa* LMG 23880<sup>T</sup> | *Ct* DSM 15264 | *Ct* LMG 21645 | *Ct* DSM 15344<sup>T</sup> |
| --- | --- | --- | --- | --- |
| Length (bp) | 3,348,564 | 4,030,985 | 3,940,444 | 3,858,501 |
| GC content (%) | 69.34 | 70.22 | 70.20 | 70.28 |
| Polycistronic operons | 637 | 774 | 753 | 745 |
| ORFs | 3,069 | 3,856 | 3,724 | 3,670 |
| CDSs | 2,976 | 3,771 | 3,627 | 3,576 |
| Pseudogenes | 37 | 25 | 37 | 33 |
| rRNA (5S, 16S, 23S) | 2, 2, 2 | 2, 2, 2 | 2, 2, 2 | 2, 2, 2 |
| tRNA | 46 | 50 | 50 | 51 |
| ncRNA | 4 | 4 | 4 | 4 |

### Functional annotation

Genes in the annotated genomes were further clustered into 26 categories according to clusters of orthologous groups (COGs). The relative abundance of COG categories in *Caldimonas /Schlegelella* species is shown in Figure 1. Although the Function unknown (S) category is still the most prevalent group in each of the studied bacteria, the improvement in COGs annotation with COGtools reduced the number of genes in this category, and the representation of the other categories increased. All of the studied *Caldimonas /Schlegelella* species show a high similarity of genome content, although minor differences between genomes are present, especially in terms of *C. thermodepolymerans* DSM 15264 and S. aquatica LMG 23380^T^. The difference exists mainly due to the already mentioned category S, where *C. thermodepolymerans* DSM 15264 is the only genome with a relative abundance of this category exceeding 10%. Further, the group Amino acid transport and metabolism (E) constitutes more than 7.5% of studied *Caldimonas /Schlegelella* genomes except *C. thermodepolymerans* DSM 15264 with nearly 7.3%. In addition, the genome of *C. thermodepolymerans* DSM 15344^T^ comprises the highest percentage of COG unknown category compared to others, even though it is a type strain bacterium. The further variations in the other COGs generate noticeable differences between *Caldimonas /Schlegelella* species (see Figure 1, supplementary Table S1). The main dissimilarity between S. aquatica LMG 23380^T^ and *C. thermodepolymerans* strains is in the relative abundance of the categories “Signal transduction mechanism” (T) and “Cell, wall/membrane/envelope biogenesis” (M), with the two comprising over 15% of S. aquatica LMG 23380^T^ genome. Even though the S group still makes up the prevalent group of the genome, the difference between group S and M, or T, is not as marked as in *C. thermodepolymerans* strains. Detailed distribution of relative abundances in the individual categories across the studied *Caldimonas /Schlegelella* species is provided in supplementary Table S1.

**Figure 1.**
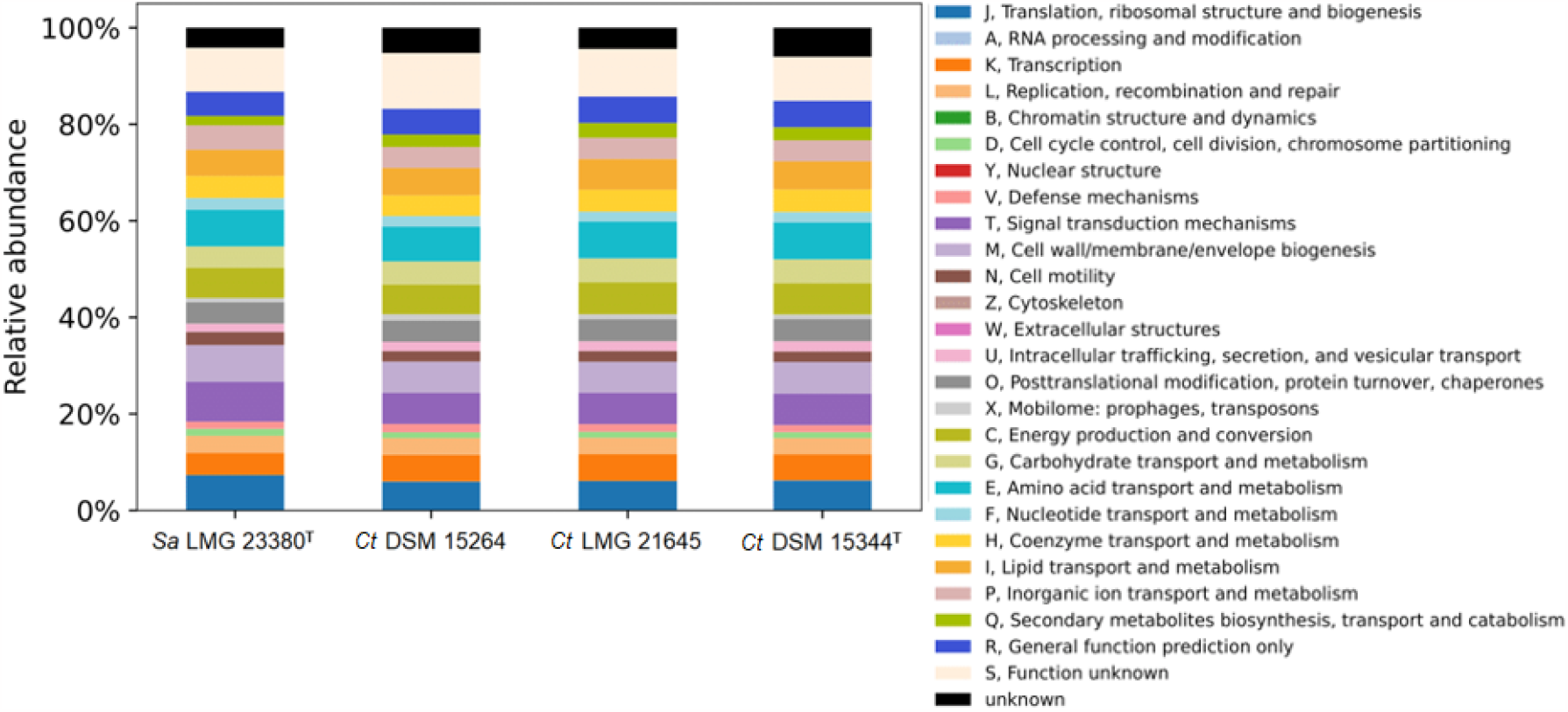
Relative abundances of genes in particular clusters of orthologous groups (COGs) categories in the *Schlegelella aquatica* (Sa) LMG 23380^T^, *Caldimonas* thermodepolymerans (Ct) DSM 15264 and LMG 21645, and reference type strain DSM 15344^T^

**Figure 2.**
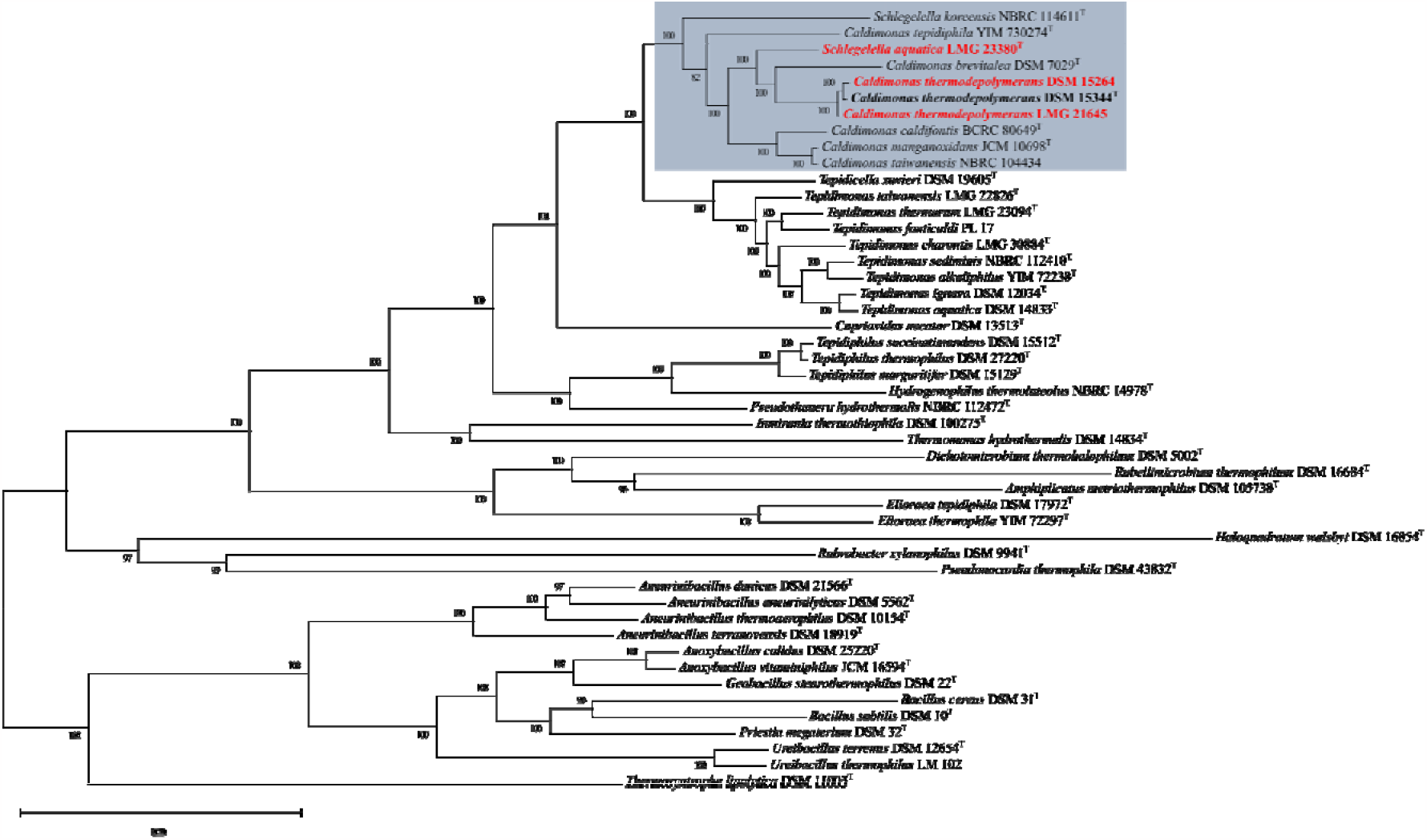
Phylogenomic tree representing placement of *Schlegelella aquatica* LMG 23880^T^ and *Caldimonas* thermodepolymerans DSM 15264 and LMG 21645 (highlighted in red). Bacteria placed in the *Caldimonas* genus are highlighted with the rectangle. The tree was constructed using the Phylophlan internal database of circa 400 genes conserved across bacterial domain. The values represent the bootstrap support based on 100 replicates.

Long ONT reads were used to detect the methylation state of cytosine nucleotides (5-mC), where a log-like ratio assigned to the target methylated site served as a measure of methylation support. The number of methylated sites in a genome differs across the studied bacteria. The highest number of methylated sites was detected in *C. thermodepolymerans* DSM 15344^T^, whereas the genome of *C. thermodepolymerans* LMG 21645 carries the lowest content of 5-mC. Subsequently, methylation motifs in bacterial genomes were derived from the previously detected methylated sites. The highest number of methylated sites in *C. thermodepolymerans D*SM 15344^T^ is consistent with its highest quantity of motifs, where six motifs satisfied the significance criteria set as 0.05 E-value. A complete overview of the methylated motifs can be found in supplementary Table S2.

Restriction-modification (R-M) analysis revealed type I, II and III systems in the studied bacteria, based on the comparison with the REBASE and additionally confirmed with the KEGG database. No complete system was found in the S. aquatica LMG 23380^T^ ; only a partial type I system was identified due to the absence of the R subunit of the system. *C. thermodepolymerans* DSM 15264 genome contains most protein-coding genes forming R-M systems, mainly genes encoding type II methylases. Furthermore, *C. thermodepolymerans* LMG 21645 possesses two complete type III R-M systems and the standalone type II methylase. The type strain of *C. thermodepolymerans* DSM 15344^T^ has three complete type I and II R-M systems. Detailed information about R-M systems of *Caldimonas /Schlegelella* species is provided in the supplementary Table S3.

Clustered Regularly Interspaced Short Palindromic Repeats (CRISPR) analysis revealed one array in each bacterium. The arrays are located near the end of each sequence, and the length of the discovered arrays ranges from 3.2kbp to 5.3kbp, except the type strain *C. thermodepolymerans* DSM 15344^T^, whose array’s length is only 163bp. In addition, one more array with the length of 1.5kbp was found in the first third of the *C. thermodepolymerans* DSM 15264 sequence. The further information is provided in the Supplementary Table S4. Cas-like genes have not been directly identified in the arrays, however, several such genes are located in their close proximity (see supplementary Table S5). Putative cas9 genes were identified in the genome of *C. thermodepolymerans* DSM 15264.

The antibiotic-resistance genes search was carried out through comparison of the sequences against the Comprehensive Antibiotic Resistance Database (CARD) and revealed three genes: *adeF*, ANT(3’’)-IIa, and aadA6 originally coding for a membrane fusion protein of the multidrug efflux complex AdeFGH *(adeF)* and aminoglycoside nucleotidyltransferases (ANT(3’’)-IIa, aadA6). The first hit, *adeF* gene, was detected in all of the *C. thermodepolymerans* strains with 71% identity of matching region and length corresponding to the criterion 100%lll± 5%. In contrast, the second gene ANT(3’’)-IIa was discovered only in the *C. thermodepolymerans* LMG 21645 strain. The aadA6 gene was identified only in the *C. thermodepolymerans* type strain DSM 15344^T^. The comparison of S. aquatica LMG 23380^T^ genome did not reveal significant hits with at least 50% identity of the matching region. Details are provided in the supplementary Table S6.

The subsequent experimental verification of susceptibility to five selected antibiotics revealed that all of the studied bacteria are sensitive to kanamycin, gentamycin, tetracycline, and chloramphenicol. The minimal inhibitory concentrations of these antibiotics determined by our probe are shown in Table 2 (the raw data can be found in the Supplementary File 2). In contrast, all three *C. thermodepolymerans* (Ct) strains grew even when the highest concentration of streptomycin (500 μg/mL) was used. Streptomycin was also the least thermally stable antibiotic in the tested set. The only exception was S. aquatica (Sa), whose growth was fully inhibited with 400 μg/mL streptomycin. Interestingly, aadA6 and ANT(3’’)-IIa genes, which can confer resistance to streptomycin, were found only in the genomes of DSM 15344^T^ and LMG 21645 strains. On the other hand, S. aquatica showed higher resistance to kanamycin and gentamycin than all of the Ct strains.

**Table 2.** Minimal inhibitory concentrations of antibiotics in NB medium. The susceptibility to the tested antibiotics is demonstrated by minimal inhibitory concentrations (MIC, the lowest concentration of antibiotic, in μg/mL, necessary to inhibit visible bacterial growth).

| Species | Kanamycin | Streptomycin | Gentamycin | Tetracycline | Chloramphenicol |
| --- | --- | --- | --- | --- | --- |
| ***Sa* LMG 23880<sup>T</sup>** | 12.5 | 400.0 | 25.0 | 5.0 | 5.0 |
| ***Ct* DSM 15264** | 2.5 | resistant | 2.5 | 12.5 | 5.0 |
| ***Ct* LMG 21645** | 7.0 | resistant | 7.0 | 5.0 | 5.0 |
| ***Ct* DSM 15344<sup>T</sup>** | 2.5 | resistant | 2.5 | 5.0 | 2.5 |

### Genome-based phylogeny and taxonomic descriptions

Digital DNA-DNA hybridization (dDDH) confirmed the classification of *C. thermodepolymerans* DSM 15264 and *C. thermodepolymerans* LMG 21645 within the *Caldimonas thermodepolymerans* species. The former exhibited 94% and the latter 95% similarity to the type strain. In contrast, S. aquatica LMG 23380^T^ exhibited a similarity of less than 30% to the *Caldimonas thermodepolymerans* species.

Phylogenomic analysis *of Caldimonas /Schlegelella* species and their comparison to the other genera of the family *Sphaerotilaceae* and additional Gram-positive and Gram-negative PHA-producing bacteria showed a well-distinguished cluster of *Caldimonas /Schlegelella* genomes confirming that these two genera are synonymous. Moreover, both *Schlegelella aquatica* and *Schlegelella koreensis* represent separate species as their genomes dDDH values compared to other *Caldimonas /Schlegelella* genomes are below the threshold for species dealineation. Therefore, we propose that *Caldimonas* aquatica comb. nov. and *Caldimonas* koreensis comb. nov. be the novel combinations for *Schlegelella aquatica* and *Schlegelella koreensis*, respectively.

### Taxonomic description of new combinations for species

Description of *Caldimonas* aquatica comb. nov.

*Caldimonas* aquatica (a.qua’ti.ca. L. fem. adj. aquatica, living in water)

Basonym: *Schlegelella aquatica* Chou et al. 2006

The description is the same as that of S. aquatica (Chou et al. 2006). Genomic, phylogenetic, and phenotypic evidence strongly support the placement of this species in the genus *Caldimonas*. The type strain is LMG 23380^T^ (= wcf1^T^ = BCRC 17557^T^). The genomic DNA G + C content is 69.34%. The GenBank accession numbers of the 16S rRNA gene and genome for the type strain are DQ417336.1 and CP110257.1, respectively.

### Description of *Caldimonas* koreensis comb. nov

*Caldimonas koreensis* (ko.re.en’sis. N.L. masc./fem. adj. koreensis, of Korea, from where the novel organisms were isolated)

Basonym: *Schlegelella koreensis* Chaudhary et al. 2022

The description is the same as that of *S. koreensis* (Chaudhary et al. 2021). Genomic, phylogenetic, and phenotypic evidence strongly support the placement of this species in the genus *Caldimonas*. The type strain is ID0723^T^ (= KCTC 72731^T^ = NBRC 114611^T^). The genomic DNA G + C content is 69.87%. The GenBank accession numbers of the 16S rRNA gene and genome for the type strain are KP326334.1 and JABWMJ01, respectively.

### PHA production on various sugars

The phenotypic investigation of various strains within the *Caldimonas /Schlegelella* genus focused on their growth and PHA production using xylose, glucose, and cellobiose as substrates. These sugars are commonly associated with lignocellulose-based resources that could potentially facilitate economically viable PHA production (Dietrich et al. 2019). Notably, the type strain *C. thermodepolymerans* DSM 15344^T^ exhibited a significant preference for xylose over glucose, which was identified as its crucial characteristic (Kourilova et al. 2020). Consequently, this attribute was assessed in other strains of *C. thermodepolymerans* as well as the closely related strain S. aquatica.

The results are presented in Table 3, indicating that *C. thermodepolymerans* achieves significantly higher PHA titers than S. aquatica when utilizing the specific substrates. Furthermore, all of the investigated strains of *C. thermodepolymerans* exhibited a substrate preference pattern similar to the type strain Ct DSM 15344^T^, with a notable preference for xylose over glucose. These findings suggest that this substrate preference is a distinct characteristic of this bacterial species. In contrast, S. aquatica displayed the opposite trend. Although glucose as a carbon substrate led to the highest yields in biomass as well as poly(3-hydroxybutyrate) (PHB) titer in this bacterium, its low PHA yields render it unsuitable for industrial-scale production.

**Table 3.** Comparison of PHB production of *Caldimonas /Schlegelella* strains utilizing different carbon substrates. Cultivation conditions – 72h, 50 °C, 180 rpm, 20 g/L; carbon source – cellobiose, glucose and xylose.

|  | *Sa* LMG 23880<sup>T</sup> |  | *Ct* DSM 15264 |  | *Ct* LMG 21645 |  | *Ct* DSM 15344<sup>T</sup> |  |
| --- | --- | --- | --- | --- | --- | --- | --- | --- |
| Substrate | CDW [g/L] | PHB [g/L] | CDW [g/L] | PHB [g/L] | CDW [g/L] | PHB [g/L] | CDW [g/L] | PHB [g/L] |
| Xylose | 0.51 ±0.01 | 0.03 ±0.01 | 6.22 ±0.11 | 4.21 ±0.02 | 2.78 ±0.01 | 1.66 ±0.11 | 5.50 ±0.07 | 3.63 ±0.01 |
| Glucose | 1.68 ±0.26 | 0.53 ±0.02 | 0.10 ±0.02 | *n.d.* | 0.09 ±0.01 | *n.d.* | 2.63 ±0.14 | 1.35 ±0.01 |
| Cellobiose | 0.19 ±0.03 | *n.d.* | 8.34 ±0.30 | 3.03 ±0.07 | 4.63 ±0.08 | 3.41 ±0.06 | 6.70 ±0.11 | 5.27 ±0.21 |

*n.d.* – not detected

An intriguing observation across all of the tested strains of *C. thermodepolymerans* was their ability to utilize cellobiose, which resulted in higher biomass growth compared to glucose or even xylose. This is surprising given that cellobiose consists of two glucose units linked by a β-1,4 glycosidic bond. It implies that the relatively low efficiency of glucose utilization in *C. thermodepolymerans* strains is more likely due to the inefficient glucose transport into bacterial cells rather than a deficiency in glucose metabolism.

Analysis of genes linked to PHA synthesis revealed a total of eight genes involved in PHA metabolism summarized in Table 4. These genes encode for enzymes involved in PHA synthesis (*phaA, phaB, phaC*) and degradation. All the tested strains harbor genes encoding for intracellular PHA depolymerase (phaZi) but S. *aqutica* does not contain genes encoding for extracellular PHA depolymerase (*phaZe*) indicating that, unlike C. *thermodepolymerans, S. aquatica* is not capable of decomposition of PHA materials in the environment. Further, the strains also carry *phaP* genes encoding for phasins – proteins covering the surface of PHA granules in the bacterial cells acting as an interface between water-rich cytoplasm and hydrophobic polymer (Mezzina and Pettinari 2016). In addition, the genomes of all of the tested strains also contain *phaR* gene, which is involved in the regulation of PHA metabolism. Finally, one additional gene related to PHA metabolism was identified solely in S. aquatica LMG 23380^T^; nevertheless, the metabolic function of the encoded protein is not known yet. Further, as is common in most PHA-producing Gram-negative bacteria, *phaC, phaA*, and *phbB* genes are clustered in the *phaC*AB operon in all of the examined *Caldimonas /Schlegelella* strains, which enables effective orchestration of PHA synthesis.

**Table 4.**
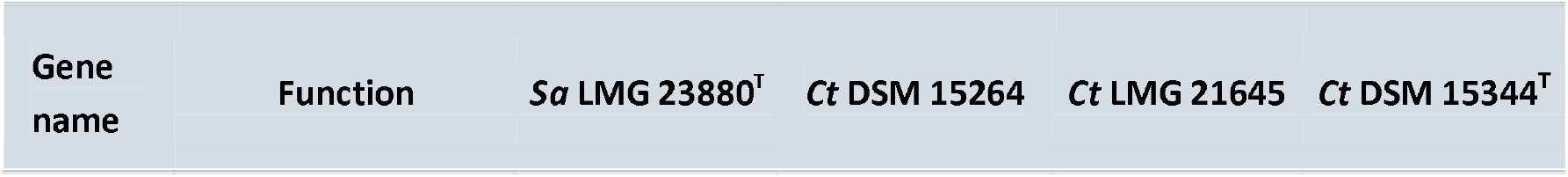
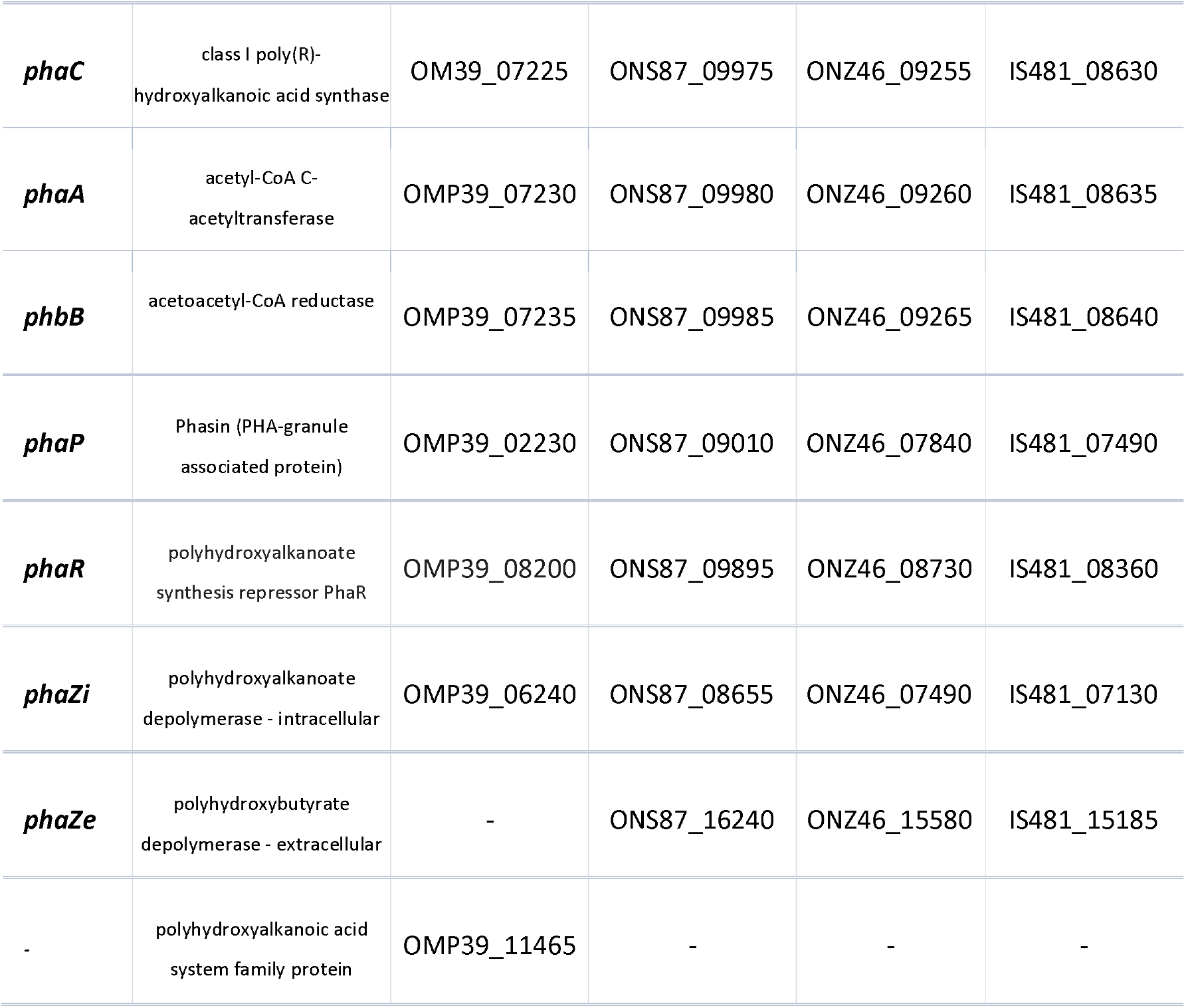
List of genes involved in polyhydroxyalkanoates metabolism in *Schlegelella aquatica* (Sa) LMG 23880^T^ *Caldimonas* thermodepolymerans (Ct) DSM 15264 and LMG 21645, and the reference type strain DSM 15344^T^, referred to by their respective locus tags. The genes forming one operon are separated from the other genes with a double line.

| Gene name | Function | *Sa* LMG 23880<sup>T</sup> | *Ct* DSM 15264 | *Ct* LMG 21645 | *Ct* DSM 15344<sup>T</sup> |
| --- | --- | --- | --- | --- | --- |
| ***phaC*** | class I poly(R)-hydroxyalkanoic acid synthase | OMP39_07225 | ONS87_09975 | ONZ46_09255 | IS481_08630 |
| ***phaA*** | acetyl-CoA C-acetyltransferase | OMP39_07230 | ONS87_09980 | ONZ46_09260 | IS481_08635 |
| ***phbB*** | acetoacetyl-CoA reductase | OMP39_07235 | ONS87_09985 | ONZ46_09265 | IS481_08640 |
| ***phaP*** | Phasin (PHA-granule associated protein) | OMP39_02230 | ONS87_09010 | ONZ46_07840 | IS481_07490 |
| ***phaR*** | polyhydroxyalkanoate synthesis repressor PhaR | OMP39_08200 | ONS87_09895 | ONZ46_08730 | IS481_08360 |
| ***phaZi*** | polyhydroxyalkanoate depolymerase - intracellular | OMP39_06240 | ONS87_08655 | ONZ46_07490 | IS481_07130 |
| ***phaZe*** | polyhydroxybutyrate depolymerase - extracellular | - | ONS87_16240 | ONZ46_15580 | IS481_15185 |
| - | polyhydroxyalkanoic acid system family protein | OMP39_11465 | - | - | - |

### Carbohydrate metabolism

To investigate the variations in the utilization of lignocellulosic sugars among the studied strains, we next focused on identifying genes responsible for encoding enzymes and transporters involved the upper carbohydrate metabolism. We confirmed the presence of genes associated with the complete Embden-Meyerhof-Parnas (EMP) pathway, which converts glucose to pyruvate and acetyl-CoA, in all four compared strains (supplementary Table S7). Regarding glucose transport into the cell, genes of the PEP-dependent phosphotransferase system (PTS) were identified in Sa LMG 23380^T^ and in all three *Caldimonas* strains, but with a fructose-specific EIICB component. Therefore, it is likely that glucose is transported by a non-PTS transporter such as the *Gts*ABCD glucose/mannose ABC transporter that we identified and whose genes form a *gts* operon in Sa LMG 23380^T^ and in *Caldimonas* strains (Figure 3). In Ct strains, the *gts* operon also includes a β-glucosidase gene and a carbohydrate porin gene. The presence of the former gene correlates with the ability of Ct strains to grow on cellobiose (Table 3).

**Figure 3.**
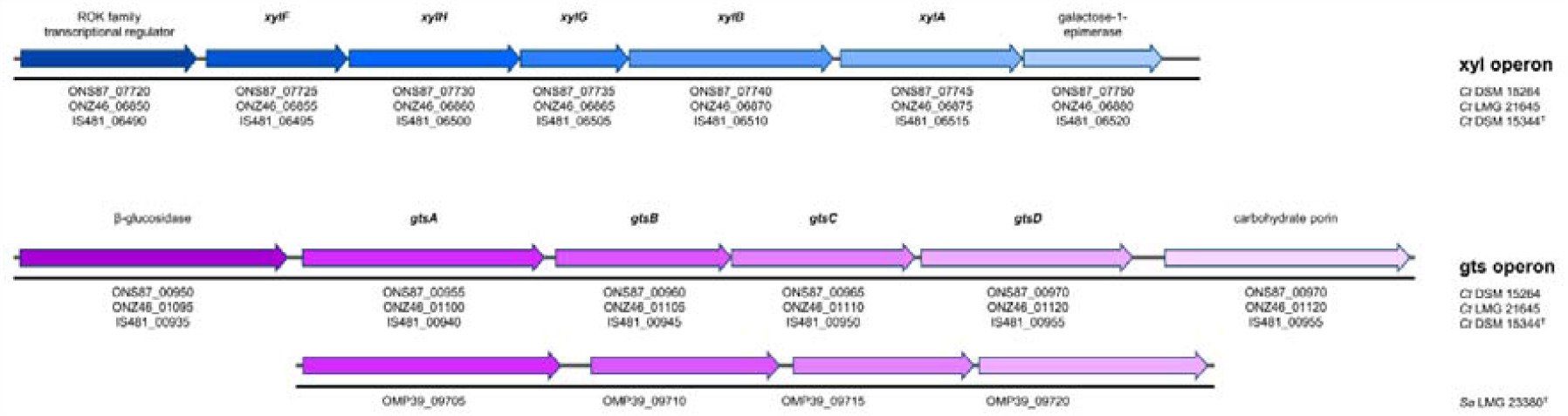
Genes involved in carbohydrate metabolism forming *xyl* and *gts* operons across *Schlegelella aquatica* (Sa) LMG 23880^T^, *Caldimonas* thermodepolymerans (Ct) DSM 15264 and LMG 21645, and the reference type strain DSM 15344^T^.

The xylose isomerase pathway (formed by xylose isomerase *XylA* and xylulose kinase *XylB*) and the non-oxidative branch of the pentose phosphate pathway (PPP) are responsible for the conversion of xylose to the EMP pathway intermediates glyceraldehyde 3-phosphate and fructose 6-phosphate in Ct DSM 15264, LMG 21645, and DSM 15344^T^ (supplementary Table S7). S. aquatica lacks *xylA* and *xylB* genes. Genes for the oxidative metabolism of xylose or glucose are not present in any of the studied strains. Interestingly, the *xylA* and *xylB* genes in Ct strains form a single *xyl* operon together with the genes encoding the high-affinity xylose ABC transporter XylFGH and a ROK-family transcriptional regulator, indicating that all *xyl* genes are regulated and expressed simultaneously (Figure 3).

An intriguing feature of the carbohydrate metabolism of all of the four compared bacteria is the absence of the two genes for enzymes of the oxidative PPP, namely 6-phosphogluconolactonase (*pg*l gene) and 6-phosphogluconate dehydrogenase (*gnd* gene). In the metabolism of the model Gram-negative bacterium Escherichia coli, *Pgl* catalyses the hydrolysis of 6-phosphogluconolactone to 6-*phosphogluconate*, which is converted by *Gnd* to ribulose 5-phosphate, which enters the non-oxidative PPP. The absence of *Gnd* is probably compensated for by the Entner-*Doudoroff* (ED) pathway formed by Edd 6-phosphogluconate dehydratase and Eda 2-keto-3-deoxy-6-*phosphogluconate* aldolase, which was found in all of the four bacteria studied and may explain their ability to grow on the lignocellulosic sugars such as xylose, glucose, and cellobiose.

## Discussion

Uncovering genomic and phenotypic traits with a focus on PHAs production by strains *Schlegelella aquatica* LMG 23380^T^ and *Caldimonas* thermodepolymerans DSM 15264 and LMG 21645 may provide important insights into microbial production of sustainable environmentally friendly polyesters. Based on our previous studies, *C. thermodepolymerans* can be considered a promising candidate for PHA production, especially from xylose-rich lignocellulose-based resources (*Kourilova* et al. 2020; *Kourilova* et al. 2021). Since the genomic information was not previously available, genome assembly was a necessary first step for further studies. De novo genome assembly using ONT reads identified one circular contig in each bacterium, indicating the absence of plasmids and thus the presence of chromosomal DNA only. Furthermore, high-quality short Illumina reads were used to polish the assemblies. The mapping of nearly all of the short reads to the final genome, as well as the prediction of the replication origin *oriC*, were unambiguous, which confirms the genome had been assembled correctly. Although genome length and GC content are higher than the average for Gram-negative bacteria (Li and Du 2014), the values are consistent with the assumption that GC content is positively correlated with genome length (Wu et al. 2012) and with growth temperature (Hu et al. 2022) as well.

Functional annotation, including the study of bacterial characteristics such as the function of individual genes, the ability to defend against foreign DNA, antibiotic resistance, and the requirements for genome editing, is the key prerequisite for understanding and manipulating the studied bacteria. Genes classified according to clusters of orthologous groups (COGs) revealed around 85% of gene functions in all of the studied *Caldimonas /Schlegelella*. However, the remaining 15% remains unknown, highlighting the uniqueness of these bacteria. Furthermore, the classification confirmed the genomic similarity among the *C. thermodepolymerans* strains and indicated a partial divergence between S. aquatica and *C. thermodepolymerans* based on different distributions across specific COGs within a narrow range. The prediction of Restriction-Modification (R-M) systems revealed the diversity of this cellular defense mechanism in the studied bacteria. The lack of active R-M systems was registered in S. aquatica LMG 23380^T^ genome, with only a partial type I system being identified, suggesting its inactivity. In contrast, all of the studied *C. thermodepolymerans* possess at least one complete R-M system, indicating a strong system protecting *C. thermodepolymerans* strains from foreign DNA, which may pose challenges in genome editing (Riley and Guss 2021). Regarding further potential opportunities for future genome engineering, all of the studied bacteria were found to have the ability to receive foreign DNA as they contain CRISPR arrays. Although these arrays serve primarily as a protection from receiving additional foreign genetic material, they can be also adopted as a reservoir of components for genome editing of *Caldimonas* strains and potentially other thermophilic bacteria (Gaj et al. 2016). Native CRISPR-Cas systems in strains S. aquatica LMG 23880^T^ and *C. thermodepolymerans* DSM 15264 and LGM 21645 will be further studied and can be repurposed for genome editing of these bacteria (Walker et al. 2020). The most promising seems CRISPR-Cas system in *C. thermodepolymerans* DSM 15264, which contains two putative cas9 genes. In contrast, the type strain *C. thermodepolymerans* DSM 15344^T^ contains arrays of very short length and no cas or cas-like genes have been identified in its chromosome, making it a suitable candidate for the adoption of exogenous CRISPR-Cas systems (Walker et al. 2020). The observed susceptibility of all *Caldimonas* strains to four out of the five selected antibiotics will facilitate their future genetic engineering as various selection markers can be used. Identified antibiotic-resistance genes can be subsequently removed to prevent a potential outbreak of antibiotic resistance.

In the genomic era, a polyphasic approach combining examinations of genotypic and phenotypic traits is still necessary and has been practiced for more than a decade in prokaryotic taxonomy (Tindall et al. 2010). Here, the classification of the studied bacteria using a combination of digital DNA-DNA hybridization (dDDH) and phylogenomic analysis confirmed that the newly analyzed non-type strains *C. thermodepolymerans* DSM 15264 and LMG 21645 belong to *C. thermodepolymerans* species and are very similar to the type strain *C. thermodepolymerans* DSM 15344^T^. Additional cultivation experiments also showed that these strains share unique phenotypic traits, particularly a preference for xylose over glucose, which seems to be a unique feature of *C. thermodepolymerans* that cannot be found in other species of the genus *Caldimonas*. This suggests, along with the fact that C. thermodepolymerans, formerly Schlegellela thermodepolymerans, was the type species of the genus Schlegelella covering four species, that the genus *Caldimonas /Schlegelella* is much more diverse than previously assumed. The previous absence of representative genomes of S. aquatica and S. koreensis, coupled with the limitations of the 16S rRNA gene in inferring the phylogeny within the family Sphaerotilaceae (Liu et al. 2022) prevented their accurate taxonomic placement. An analysis of the newly assembled S. aquatica and publicly available S. koreensis genomes revealed that *Schlegelella aquatica* and *Schlegelella koreensis* are homotypic synonyms for *Caldimonas aquatica* and *Caldimonas koreensis*, respectively. At the same time, genotypic analysis proved that both organisms represent separate species, as also shown by their different phenotypes. S. aquatica, in comparison to *C. thermodepolymerans*, exhibits slower growth and lower production of PHAs when utilizing various carbon sources and prefers glucose over xylose. Although we did not analyze the phenotype of *S. koreensis*, this species had been already shown to possess unique traits such as different temperature ranges for growth and the inability to utilize various saccharides while still producing PHA granules (Chaudhary et al. 2021).

PHAs can be synthesized through four different pathways (Steinbüchel 1991). Primarily, a three-step pathway, encoded by the genes forming the *phaC*AB operon (Umeda et al. 1998; Kourilova et al. 2020), synthetizes PHAs from acetyl-CoA. This key metabolic intermediate is converted into acetoacetyl-CoA, followed by the conversion into 3-hydroxybutyryl-CoA (Naik et al. 2008; Pradhan et al. 2020). Given that the *phaC*AB operon has been identified in each of the studied *Caldimonas* strains, exploitation of this pathway during the growth on the assessed lignocellulosic sugars (xylose, glucose, and cellobiose) can be assumed (Figure 4). Additional pathways of PHAs metabolism include carbohydrate biosynthesis, fatty acid de novo synthesis, and fatty acid β-oxidation. Genes involved in these three pathways have been identified in all the bacteria studied, which likely implies the ability to synthesize PHAs by all the pathways mentioned; however, further research is needed to confirm this hypothesis. In addition, several differences between S. aquatica (Sa) and *C. thermodepolymerans* (Ct) have been identified. Firstly, S. aquatica most likely does not possess the enzymatic apparatus for extracellular degradation of PHA materials present in the environment. On the other hand, *C. thermodepolymerans* was found to possess highly effective extracellular PHA depolymerase. In fact, the great PHA degradation efficiency is one of the most notable features of C. thermodepolymerans, which is also reflected in its taxonomic name: thermodepolymerans – i.e. capable of degradation of PHA polymers under thermophilic conditions (Elbanna et al. 2003). Further, a gene belonging to the polyhydroxyalkanoic acid group family with unknown function is exclusively present in the genome of Sa.

**Figure 4.**
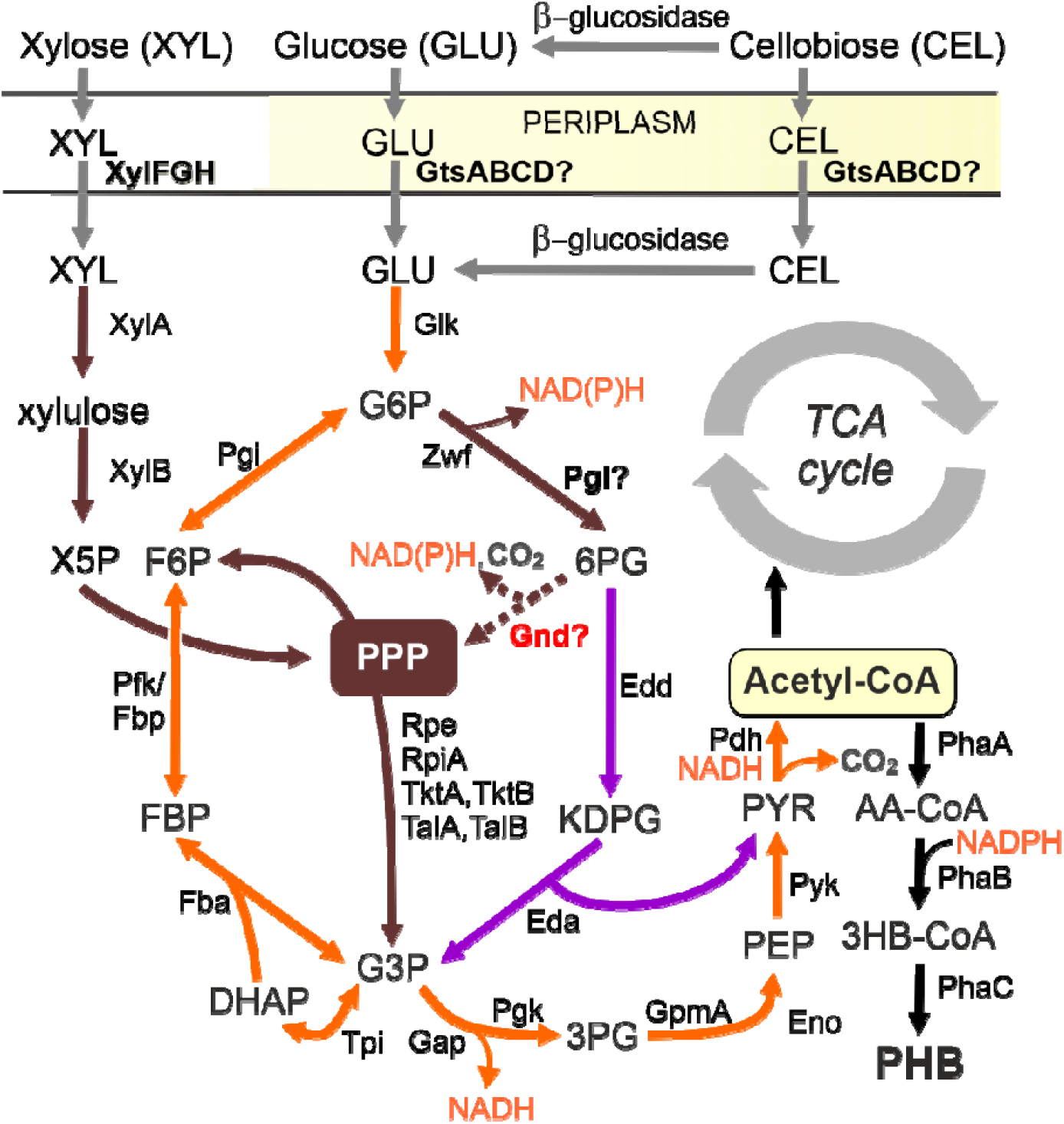
Schematic illustration of proposed carbohydrate metabolism in studied *Caldimonas /Schlegelella* strains. Xylose isomerase pathway (formed by *XylA* xylose isomerase and *XylB* xylulokinase) and the pentose phosphate pathway are shown using brown arrows, the Embden-Meyerhof-Parnas pathway is shown with orange arrows, and the Entner-Doudoroff pathway with magenta arrows. Abbreviations (enzymes): Eda, 2-keto-3-deoxy-6-phosphogluconate aldolase; Edd, 6-phosphogluconate dehydratase; Eno, phosphopyruvate hydratase; Fba, fructose-1,6-biphosphate aldolase; Fbp, fructose-1,6-biphosphatase; Gap, glyceraldehyde-3-phosphate dehydrogenase; Gcd, glucose dehydrogenase; Gnd, 6-phosphogluconate dehydrogenase; Pdh, pyruvate dehydrogenase; Pgi, glucose-6-phosphate isomerase; Pgk, phosphoglycerate kinase; Pgl, 6-phosphogluconolactonase; Pgm, phosphoglycerate mutase; *PhaA*, acetyl-CoA acetyltransferase; *PhaB*, acetoacetyl-CoA reductase; *PhaC*, poly(3-hydroxyalkanoate) polymerase; Pyk, pyruvate kinase; Rpe, ribulose-5-phosphate 3-epimerase; RpiA, ribose-5-phosphate isomerase; Tal, transaldolase; Tkt, transketolase; Tpi, triosephosphate isomerase; Zwf glucose-6-phosphate dehydrogenase. Abbreviations (metabolites): AA-CoA, acetoacetyl coenzyme A; Acetyl-CoA, acetyl coenzyme A; DHPA, dihydroxyacetone phosphate; E4P, erythrose 4-phosphate; FBP, fructose 1,6-biphosphate; F6P, fructose 6-phosphate; GLL, glucono-δ-lactone; GLN, gluconate; G3P glyceraldehyde 3-phosphate; G6P, glucose 6-phosphate; 3HB-CoA, 3-hydroxybutyryl coenzyme A; KDPG, 2-keto-3-deoxy-6-phosphogluconate; 2KG, 2-ketogluconate; 2KG-6P, 2-ketogluconate 6-phosphate; NADH, reduced nicotinamide adenine dinucleotide; NADPH, reduced nicotinamide adenine dinucleotide phosphate; PEP, phosphoenolpyruvate; 3PG, 3-phosphoglycerate; 6PG, 6-phosphogluconate; PHB, poly-3-hydroxybutyrate; R5P, ribose 5-phosphate; Ru5P, ribulose 5-phosphate; S7P, seduheptulose 7-phosphate; XLN, xylonate; X5P, xylulose 5-phosphate. *Gts*ABCD and XylFGH stand for mannose/glucose and xylose ABC transporter, respectively. Note that xylose isomerase pathway and b-glucosidase are not present in S. aquatica.

Genomic differences were also confirmed by the cultivation experiments. The ability to produce PHA was demonstrated in all of the investigated strains. However, a significantly lower ability to utilize carbohydrate substrates and produce PHA was observed in S. aquatica LMG 23380^T^. *C. thermodepolymerans* DSM 15264 showed the greatest similarity to the type strain *C. thermodepolymerans* DSM 15344^T^. A high increase in biomass was observed for both cellobiose and xylose substrates. The contrasting lack of growth and PHA formation in S. aquatica on xylose and cellobiose might be explained by the absence of the *xyl* operon and the β-glucosidase gene in its genome. Unlike S. aquatica, all Ct strains possess these genes, as reflected in their ability to utilize xylose and cellobiose. It is not clear now why all of the Ct strains grow much worse on glucose than on cellobiose. Since all the bacteria investigated in this study possess genes assumed to encode metabolic traits that ensure glucose utilization (Figure 4), we hypothesize that the bottleneck limiting the utilization of this sugar may lie in its transport across the cytoplasmic membrane. The identified *Gts*ABCD glucose/mannose transporter might have a higher affinity for cellobiose than for monomeric glucose, whose transport would be either slow (Sa LMG 23380^T^, Ct DSM 15344^T^) or negligible (Ct DSM 15264 and Ct LMG 21645). Uptake of cellobiose and other cellooligosaccharides through the ABC-type glucose transporters is common in bacteria (Parisutham et al. 2017; Dvořák and de Lorenzo 2018). The transport of oligosaccharides is more economical with regards to ATP than glucose transport. Our hypothesis is further supported by the presence of the β-glucosidase gene in the *gts* operon of Ct strains and by the fact that residual glucose is often detected in culture supernatants of the strains grown on cellobiose (data not shown). The latter observation also implies that at least a part of the cellobiose substrate is cleaved to glucose monomers outside the cell. A signal sequence specific for the twin-arginine translocation (Tat) pathway was identified at the 5’ end of the β-glucosidase genes in the Ct strains. Since the Tat pathway is known to translocate already folded (and active) proteins across the cytoplasmic membrane (Berks et al. 2003), we argue that cellobiose hydrolysis by β-glucosidase in the Ct strains studied takes place both in the cytoplasm and outside the cell (Figure 4).

The proposed scheme of the upper carbohydrate metabolism in the studied thermophilic PHA producers (Figure 4) suggests that all three tested lignocellulosic sugars can be utilized, despite the lack of pgl and gnd homologs in the *Caldimonas* genomes. Hydrolysis of 6-phosphogluconolactone can occur spontaneously (Fatima et al. 2018), although it has been suggested that Pgl lactonase accelerates the reaction and prevents the δ form of the lactone from being converted to the dead-end γ form (Miclet et al. 2001). A possible compensatory mechanism for the lack of Gnd is gluconate 6-phosphate undergoing the ED shunt to glyceraldehyde 3-phosphate and pyruvate. This way, 6-*phosphogluconate* would not become a dead-end product and at least 1 mol of NADPH per 1 mol of glucose would be produced in the glucose-6-phosphate dehydrogenase (Zwf) reaction in the oxidative branch of the PPP. During the growth on glucose or cellobiose, at least a small reverse flux from glyceraldehyde 3-phosphate and fructose 6-phosphate to the non-oxidative PPP is required to ensure the formation of nucleotide precursors. Hence, one the preference of xylose over glucose in *Caldimonas* strains might be explained by the fact that the pathway for securing important metabolic precursors in PPP such as ribulose 5-phosphate is shorter with pentose. However, more experimental data is necessary to investigate this phenomenon.

In conclusion, the genomic and phenotypic characterization of the non-model bacteria *Schlegelella aquatica* LMG 23380^T^ and *Caldimonas* thermodepolymerans DSM 15264 and LMG 21645 provides valuable insights into microbial production of polyhydroxyalkanoates, sustainable and environmentally friendly polyesters. The genome assembly and functional annotation of these bacteria confirm their potential with regards to PHA production and reveal their individual characteristics. Particularly, the unique *xyl* operon present only in *C. thermodepolymerans* suggests their strong potential for biotechnological PHA production from xylose-rich resources. Furthermore, the fact that the strains of *C. thermodepolymerans* are also capable of utilizing cellobiose is of biotechnological interest since cellobiose is commonly present in lignocellulose-based media when enzymatic hydrolysis of cellulose is employed. Further, the fact that the studied strains of *C. thermodepolymerans* are capable of cellobiose utilization also indicates that the low efficiency of glucose metabolism might be linked to difficulties with the transport of glucose into the cells rather than a deficiency in its metabolization in the cells. In this case, glucose uptake and conversion into PHA might be potentially improved by introducing heterologous powerful glucose transporter using approaches of synthetic biology. Therefore, further investigation is required to fully understand the organism’s properties and improve its potential for PHA production in agreement with the concept of NGIB.

## Acknowledgments

This study was supported by the grant project GACR GA22-10845S and from the European Union’s Horizon 2020 research and innovation programme under the Marie Sklodowska-Curie grant agreement No. 101023766. PhD study of Jana Musilova is supported by the Brno Ph.D. Talent Scholarship – Funded by the Brno City Municipality.

## Data availability

The whole-genome sequences of S. aquatica LMG 23380^T^, *C. thermodepolymerans* DSM 15264 and *C. thermodepolymerans* LMG 21645 have been deposited in the DDBJ/ENA/GenBank under the accessions CP110257, CP110416, and CP110415, respectively. The raw paired-end Illumina reads have been deposited in the NCBI SRA database under the accession numbers SRX20788156, SRX20788022, and SRX20788024; Oxford Nanopore Technologies raw reads are accessible under the numbers SRX20788157, SRX20788023, and SRX20788025.

## Author contributions

Jana *Musilova*: Methodology, Software, Formal analysis, Investigation, Data Curation, Writing - Original Draft, Visualization. *Xenie Kourilova*: Methodology, Validation, Formal analysis, Investigation, Writing - Original Draft. Kristyna *Hermankova*: Software, Formal analysis, Data Curation, Writing - Original Draft, Visualization. Matej Bezdicek: Investigation, Writing - Original Draft. Anastasiia Ieremenko: Investigation. Pavel Dvorak: Conceptualization, Formal analysis, Writing - Original Draft, Visualization, Funding acquisition. *Stanislav Obruca*: Conceptualization, Writing - Original Draft, Visualization, Funding acquisition. Karel Sedlar: Conceptualization, Formal analysis, Data Curation, Writing - Original Draft, Visualization, Funding acquisition.

