## Supplementary File 1 for "Genomic and Phenotypic Comparison of Polyhydroxyalkanoates Producing Strains of genus *Caldimonas/Schlegelella*"

### Supplements

**Table S1**

Clusters of orthologous groups (COGs) categories in the *Schlegelella aquatica* (*Sa*) LMG 23380$^{T}$, *Caldimonas thermodepolymerans* (*St*) strains DSM 15264, LMG 21645, and reference type strain DSM 15344$^{T}$.

| Category | *Sa* LMG 23380$^{T}$ [%] | *Ct* DSM 15264 [%] | *Ct* LMG 21645 [%] | *Ct* DSM 15344$^{T}$ [%] |
|---|---|---|---|---|
| J, Translation, ribosomal structure and biogenesis | 7.30 | 5.97 | 6.08 | 6.20 |
| A, RNA processing and modification | 0.03 | 0.03 | 0.03 | 0.03 |
| K, Transcription | 4.61 | 5.55 | 5.56 | 5.39 |
| L, Replication, recombination and repair | 3.52 | 3.39 | 3.38 | 3.34 |
| B, Chromatin structure and dynamics | 0.03 | 0.03 | 0.03 | 0.03 |
| D, Cell cycle control, cell division, chromosome partitioning | 1.43 | 1.21 | 1.23 | 1.22 |
| V, Defense mechanisms | 1.49 | 1.74 | 1.58 | 1.47 |
| T, Signal transduction mechanisms | 8.29 | 6.42 | 6.54 | 6.56 |
| M, Cell wall/membrane/envelope biogenesis | 7.56 | 6.50 | 6.38 | 6.51 |
| N, Cell motility | 2.72 | 2.16 | 2.21 | 2.17 |
| Z, Cytoskeleton | 0.00 | 0.03 | 0.03 | 0.03 |
| W, Extracellular structures | 0.00 | 0.03 | 0.03 | 0.03 |
| U, Intracellular trafficking, secretion, and vesicular transport | 1.72 | 1.89 | 1.99 | 2.09 |
| O, Posttranslational modification, protein turnover, chaperones | 4.44 | 4.45 | 4.55 | 4.56 |
| X, Mobilome: prophages, transposons | 0.86 | 1.26 | 1.04 | 0.97 |
| C, Energy production and conversion | 6.27 | 6.18 | 6.62 | 6.48 |
| G, Carbohydrate transport and metabolism | 4.41 | 4.76 | 4.96 | 4.95 |
| E, Amino acid transport and metabolism | 7.63 | 7.23 | 7.58 | 7.67 |
| F, Nucleotide transport and metabolism | 2.39 | 2.16 | 2.10 | 2.14 |
| H, Coenzyme transport and metabolism | 4.61 | 4.34 | 4.52 | 4.70 |
| I, Lipid transport and metabolism | 5.41 | 5.60 | 6.38 | 5.81 |
| P, Inorganic Ion transport and metabolism | 5.04 | 4.37 | 4.42 | 4.34 |
| Q, Secondary metabolites biosynthesis, transport and catabolism | 1.96 | 2.58 | 3.05 | 2.70 |
| R, General function prediction only | 5.01 | 5.34 | 5.51 | 5.48 |
| S, Function unknown | 9.12 | 11.55 | 9.84 | 9.12 |
| COG unknown | 4.15 | 5.24 | 4.39 | 6.03 |

**Table S2**

Methylation motifs detected in the *Schlegelella aquatica* LMG 23380$^T$, *Caldimonas thermodepolymerans* strains DSM 15264, LMG 21645, and reference type strain DSM 15344$^T$.

| Motif | E-value | Methylated sites count |
|---|---|---|
| ***Schlegelella aquatica* LMG 23380$^T$** | | |
| WGGNCGASMHSSNSS | 2.0e-007 | 25,498 |
| NWGSCGSWGN | 3.9e-006 | |
| TCCTCGNYCW | 6.7e-005 | |
| ***Caldimonas thermodepolymerans* DSM 15264** | | |
| NWSBCGVTGNYSSNS | 5.9e-024 | 46,563 |
| WSCTCGDCSHSGNSS | 1.6e-021 | |
| SCWGCGCCWG | 1.0e-011 | |
| WGGRCGDSRW | 1.5e-007 | |
| CTCGCTCA | 8.3e-003 | |
| ***Caldimonas thermodepolymerans* LMG 21645** | | |
| BCWGCGSSWSS | 1.3e-002 | 12,898 |
| WGGNCGRGVW | 5.5e-002 | |
| ***Caldimonas thermodepolymerans* DSM 15344$^T$** | | |
| VWSSCGSTGVYSSNS | 4.0e-064 | 67,944 |
| WGSYCGRCCWSSNCG | 1.9e-031 | |
| SAGGCGCTGV | 1.9e-010 | |
| WSSYCGSCGRSSW | 1.8e-003 | |
| CCAGCGTGCCGNCGM | 1.2e-002 | |
| WNNHCGDNNW | 1.3e-002 | |

**Table S3**

Restriction-Modification (R-M) systems detected in the *Schlegelella aquatica* LMG 23380$^{T}$, *Caldimonas thermodepolymerans* strains DSM 15264, LMG 21645, and reference type strain DSM 15344$^{T}$. Prediction by REBASE database (normal text) is enriched by the match with KEGG database prediction (bold text).

| **R-M systems type** | | | | | | |
|---|---|---|---|---|---|---|
| **I** | | | **II** | | **III** | |
| **Gene**[a] | | | | | | |
| **R** | **S** | **M** | **R** | **M** | **R** | **M** |
| ***Schlegelella aquatica* LMG 23380$^{T}$** | | | | | | |
| - | **OMP39_01035** | **OMP39_01025** | - | - | - | - |
| ***Caldimonas thermodepolymerans* DSM 15264** | | | | | | |
| **ONS87_07805** | **ONS87_07810** | **ONS87_07820** | - | ONS87_02285<br>ONS87_02300 | - | - |
| **ONS87_15895** | | | - | ONS87_02485 | - | - |
| ONS87_15865 | **ONS87_15850** | **ONS87_15840** | - | ONS87_07330<br>ONS87_07335 | - | - |
| ONS87_15875 | | | - | ONS87_07990<br>ONS87_07995 | - | - |
| - | - | - | ONS87_15885 | ONS87_15880 | - | - |
| ***Caldimonas thermodepolymerans* LMG 21645** | | | | | | |
| - | - | - | - | ONZ46_6505<br>ONZ46_6510 | **ONZ46_06670** | **ONZ46_06675** |
| - | - | - | - | - | ONZ46_09195 | **ONZ46_09205** |
| ***Caldimonas thermodepolymerans* DSM 15344$^{T}$** | | | | | | |
| **IS481_08585** | **IS481_08575** | **IS481_08580** | IS481_14855 | **IS481_14860** | - | - |
| - | - | - | IS481_14025 | IS481_14020 | - | - |

[a] *R* restriction endonuclease, *M* restriction endonuclease coupled methylation protein, *S* R-M specific protein

**Table S4**

Antibiotic resistance genes in the *Schlegelella aquatica* LMG 23380ᵀ, *Caldimonas thermodepolymerans* strains DSM 15264, LMG 21645, and reference type strain DSM 15344ᵀ.

| Antibiotic Resistance Ontology (ARO) term | Drug class | Resistance mechanism | AMR Gene Family | % identity of matching region |
|---|---|---|---|---|
| ***Schlegelella aquatica* LMG 23380ᵀ** | | | | |
| adeF | fluoroquinolone antibiotic, tetracycline antibiotic | antibiotic efflux | resistance-nodulation-cell division (RND) antibiotic efflux pump | 46.24 |
| ***Caldimonas thermodepolymerans* DSM 15264** | | | | |
| adeF | fluoroquinolone antibiotic, tetracycline antibiotic | antibiotic efflux | resistance-nodulation-cell division (RND) antibiotic efflux pump | 46.74 |
| adeF | fluoroquinolone antibiotic, tetracycline antibiotic | antibiotic efflux | resistance-nodulation-cell division (RND) antibiotic efflux pump | 71.54 |
| ***Caldimonas thermodepolymerans* LMG 21645** | | | | |
| adeF | fluoroquinolone antibiotic, tetracycline antibiotic | antibiotic efflux | resistance-nodulation-cell division (RND) antibiotic efflux pump | 46.74 |
| adeF | fluoroquinolone antibiotic, tetracycline antibiotic | antibiotic efflux | resistance-nodulation-cell division (RND) antibiotic efflux pump | 71.44 |
| ANT(3'')-IIa | ANT(3'') | antibiotic inactivation | aminoglycoside antibiotic | 98.52 |
| ***Caldimonas thermodepolymerans* DSM 15344ᵀ** | | | | |
| aadA6 | aminoglycoside antibiotic | antibiotic inactivation | aminoglycoside antibiotic | 100.00 |
| adeF | fluoroquinolone antibiotic, tetracycline antibiotic | antibiotic efflux | resistance-nodulation-cell division (RND) antibiotic efflux pump | 46.74 |
| adeF | fluoroquinolone antibiotic, tetracycline antibiotic | antibiotic efflux | resistance-nodulation-cell division (RND) antibiotic efflux pump | 71.25 |

**Table S5**

CRISPR arrays in the *Schlegelella aquatica* LMG 23380$^T$, *Caldimonas thermodepolymerans* strains DSM 15264, LMG 21645, and reference type strain DSM 15344$^T$.

| Start | End | Score | Representative repeat (RR) | RR length | No. of spacers |
|---|---|---|---|---|---|
| ***Schlegelella aquatica* LMG 23380$^T$** | | | | | |
| 3,020,369 | 3,023,539 | 7.94 | GTTCGCTGCCGCG TAGGCAGCTCAGA AA | 28 | 52 |
| ***Caldimonas thermodepolymerans* DSM 15264** | | | | | |
| 653,366 | 654,948 | 7.29 | GAGTGTAGCTATC CGGGGTGAGAGA GGAAGCTACAAC | 37 | 23 |
| 3,620,825 | 3,626,100 | 7.79 | GTCTTCCCCGCGT GAGCGGGGATCG ACCC | 29 | 86 |
| ***Caldimonas thermodepolymerans* LMG 21645** | | | | | |
| 3,529,982 | 3,533,425 | 7.44 | GTCTTCCCCGCGT GAGCGGGGATCG ACC | 28 | 56 |
| ***Caldimonas thermodepolymerans* DSM 15344$^T$** | | | | | |
| 3,438,037 | 3,438,200 | 5.05 | GTCGCGCCCTCAC GGGCGCGTGGGT TGAAAC | 31 | 2 |

**Table S6**

Cas-like genes identified in the *Schlegelella aquatica* (*Sa*) LMG 23380$^T$, *Caldimonas thermodepolymerans* (*St*) strains DSM 15264, LMG 21645, and reference type strain DSM 15344$^T$; referred by individual bacterium´s locus tag and its position in genome.

| Gene name | *Sa* LMG 23380$^T$ | *Ct* DSM 15264 | *Ct* LMG 21645 | *Ct* DSM 15344$^T$ |
|---|---|---|---|---|
| *cas1f* | OMP39_13840<br>3,010,335 – 3,011,312 | - | - | - |
| *cas3f* | OMP39_13845<br>3,011,309 – 3,014,707 | - | - | - |
| *cas6f* | OMP39_13870<br>3,019,672 – 3,020,238 | - | - | - |
| *cas2* | - | ONS87_03305<br>655,010 – 655,318 | - | - |
| *cas1* | - | ONS87_03310<br>655,354 – 656,259 | - | - |
| *cas9* | - | ONS87_03315<br>656,246 – 659,608 | - | - |
| *cas3* | - | ONS87_17330<br>3,612,301 – 3,614,844 | ONZ46_16665<br>3,521,458 – 3,524,001 | - |
| *casA* | - | ONS87_17335<br>3,614,837 – 3,616,447 | ONZ46_16670<br>3,523,994 – 3,525,604 | - |
| *casB* | - | ONS87_17340<br>3,616,444 – 3,616,971 | ONZ46_16675<br>3,525,601 – 3,526,128 | - |
| *cas7e* | - | ONS87_17345<br>3,616,977 – 3,618,191 | ONZ46_16680<br>3,526,134 – 3,527,348 | - |
| *cas5e* | - | ONS87_17350<br>3,618,195 – 3,168,908 | ONZ46_16685<br>3,527,352 – 3,528,065 | - |
| *cas1e* | - | ONS87_17360<br>3,619,600 – 3,620,499 | ONZ46_16695<br>3,528,757 – 3,529,656 | - |
| *cas2e* | - | ONS87_17365<br>3,620,471 – 3,620,782 | ONZ46_16700<br>3,529,628 – 3,529,939 | - |

**Table S7**

Components of carbohydrate metabolism in the *Schlegelella aquatica* (*Sa*) LMG 23380$^T$, *Caldimonas thermodepolymerans* (*St*) strains DSM 15264, LMG 21645, and reference type strain DSM 15344$^T$; referred by individual bacterium´s locus tag.

| Component | *Sa* LMG 23380$^T$ | *Ct* DSM 15264 | *Ct* LMG 21645 | *Ct* DSM 15344$^T$ |
|---|---|---|---|---|
| **Glucose metabolism** | | | | |
| **GtsABCD** Glucose/manose ABC transporter (glucose-binding periplasmic protein, permease protein, permease, ATP binding protein) | OMP39_09705<br>OMP39_09710<br>OMP39_09715<br>OMP39_09720 | ONS87_00955<br>ONS87_00960<br>ONS87_00965<br>ONS87_00970 | ONZ46_01100<br>ONZ46_01105<br>ONZ46_01110<br>ONZ46_01115 | IS481_00940<br>IS481_00945<br>IS481_00950<br>IS481_00955 |
| **Embden-Meyerhof-Parnas pathway** | | | | |
| **Glk** Glucokinase | OMP39_00720<br>OMP39_07045 | ONS87_01110 | ONZ46_01255 | IS481_01095 |
| **Pgi** Glucose-6-phosphate isomerase | OMP39_11490 | ONS87_04135 | ONZ46_03730 | IS481_03595 |
| **PfkA** ATP-dependent 6-phosphofructokinase | OMP39_12170 | ONS87_13450 | ONZ46_12795 | IS481_12325 |
| **Fbp** Fructose-1,6-bisphosphatase | OMP39_04185 | ONS87_05860 | ONZ46_05415 | IS481_05335 |
| **FbaA** Fructose 1,6-biphosphate aldolase | OMP39_11720 | ONS87_03905 | ONZ46_03510 | IS481_03365 |
| **TpiA** Triose phosphate isomerase | OMP39_09860 | ONS87_12695 | ONZ46_12045 | IS481_11345 |
| **GapA** Glyceraldehyde-3-phosphate dehydrogenase | OMP39_01410 | ONS87_02020 | ONZ46_02100 | IS481_01955 |
| **Pgk** Phosphoglycerate kinase | OMP39_11710 | ONS87_03920 | ONZ46_03525 | IS481_03380 |
| **GpmA** 2,3-bisphosphoglycerate-dependent phosphoglycerate mutase | OMP39_02970 | ONS87_15510 | ONZ46_14875 | IS481_14385 |
| **Eno** phosphopyruvate hydratase/enolase | OMP39_06045 | ONS87_08465 | ONZ46_07300 | IS481_06940 |
| **Pyk** Pyruvate kinase I and II | OMP39_11715 | ONS87_03915 | ONZ46_03520 | IS481_03375 |
| **Pdh** Pyruvate dehydrogenase complex (pyruvate dehydrogenase E1 component AceE, dihydrolipoyllysine-residue acetyltransferase E2 component AceF, dihydrolipoyl dehydrogenase E3 component LpdA) | OMP39_06485<br>OMP39_06490<br>OMP39_06495 | ONS87_08915<br>ONS87_08920<br>ONS87_08925 | ONZ46_07745<br>ONZ46_07750<br>ONZ46_07755 | IS481_07395<br>IS481_07400<br>IS481_07405 |
| **Entner-Doudoroff pathway** | | | | |
| **Edd** 6-phosphogluconate dehydratase | OMP39_10305 | ONS87_13820 | ONZ46_13170 | IS481_12690 |
| **Eda** 2-Keto-3-deoxy-6-phosphogluconate aldolase | OMP39_10300 | ONS87_13815 | ONZ46_13165 | IS481_12685 |
| **Cellobiose metabolism** | | | | |
| β-glucosidase | - | ONS87_00950 | ONZ46_01095 | IS481_00935 |
| **Xylose metabolism** | | | | |
| **Xylose isomerase pathway** | | | | |
| **XylFGH** xylose ABC transporter (xylF xylose-binding periplasmic protein, xylH permease protein, xylG ATP-binding protein) | - | ONS87_07725<br>ONS87_07730<br>ONS87_07735 | ONZ46_06855<br>ONZ46_06860<br>ONZ46_06865 | IS481_06495<br>IS481_06500<br>IS481_06505 |
| **XylB** Xylulose kinase | - | ONS87_07740 | ONZ46_06870 | IS481_06510 |
| **XylA** Xylose isomerase | - | ONS87_07745 | ONZ46_06875 | IS481_06515 |
| **Pentose phosphate pathway** | | | | |
| **Rpe** Ribulose-phosphate 3-epimerase | OMP39_13725 | ONS87_17200 | ONZ46_16535 | IS481_16145 |

| | | | | |
|---|---|---|---|---|
| **RpiA** Ribose-5-phosphate isomerase A | OMP39_06455 | ONS87_08885 | ONZ46_07715 | IS481_07365 |
| **Tkt** Transketolase (TktA, TktB) | OMP39_01405 | ONS87_02015 | ONZ46_02095 | IS481_01950 |
| **Tal** Transalsolase (TalA, TalB) | OMP39_11495 | ONS87_04130 | ONZ46_03725 | IS481_03590 |
| **Zwf** Glucose-6-phosphate 1-dehydrogenase | OMP39_11500 | ONS87_04120 | ONZ46_03720 | IS481_03585 |
